# Potential inhibitors for 2019-nCoV coronavirus M protease from clinically approved medicines

**DOI:** 10.1101/2020.01.29.924100

**Authors:** Xin Liu, Xiu-Jie Wang

## Abstract

Starting from December 2019, a novel coronavirus, named 2019-nCoV, was found to cause Severe Acute Respiratory (SARI) symptoms and rapid pandemic in China. With the hope to identify candidate drugs for 2019-nCoV, we adopted a computational approach to screen for available commercial medicines which may function as inhibitors for the M^pro^ of 2019-nCoV. Up to 10 commercial medicines that may form hydrogen bounds to key residues within the binding pocket of 2019-nCoV M^pro^ were identified, which may have higher mutation tolerance than lopinavir/ritonavir and may also function as inhibitors for other coronaviruses with similar M^pro^ binding sites and pocket structures.

## Introduction

Coronaviruses, members of the family *Coronaviridae* and subfamily *Coronavirinae*, are enveloped positive-stranded RNA viruses which have spikes of glycoproteins projecting from their viral envelopes, thus exhibit a corona or halo-like appearance (Masters and Perlman, 2013; Cui et al., 2019) Coronaviruses are the causal pathogens for a wide spectrum of respiratory and gastrointestinal diseases in both wild and domestic animals, including birds, pigs, rodents, etc (Dhama et al., 2014). Previous studies have found that six strains of coronaviruses are capable to infect human, including four strains circulating yearly to cause common cold, and other two strains which the source for severe acute respiratory syndrome (SARS) and Middle East respiratory syndrome (MERS-CoV), respectively (Cui et al., 2019; Dhama et al., 2014).

Starting from December 2019, a novel coronavirus, which was later named 2019-nCoV (‘n’ stands for novel), was found to cause Severe Acute Respiratory (SARI) symptoms, including fever, dyspnea, asthenia and pneumonia among people in Wuhan, China (Zhu et al., 2020; Lu et al., 2020; Hui et al., 2020). The first batch of patients infected by 2019-nCoV were almost all connected to a seafood market in Wuhan, which also trades wild animals. Later, contact transmission of 2019-nCoV among humans was confirmed, and the number of infected patients increased rapidly in Wuhan as well as other major cities in China. A series of actions have taken by the Chinese government to control the pandemic of the virus, and effective medical methods are in urgent needs to prevent 2019-nCoV infection and cure the disease.

Among all know RNA viruses, coronaviruses have the largest genomes ranging from 26 to 32 kb in length (Regenmortel et al., 2000; Schoeman and Fielding, 2019). Besides encoding structural proteins, majority part of the coronavirus genome is transcribed and translated into a polypeptide, which encodes proteins essential for viral replication and gene expression (Lai and Holmes, 2001). The ~306 aa long main protease (M^pro^), a key enzyme for coronavirus replication, is also encoded by the polypeptide and responsible for processing the polypeptide into functional proteins (Lai and Holmes, 2001). The M^pro^ has similar cleavage-site specificity to that of picornavirus 3C protease (3C^pro^), thus is also known as 3C-like protease (3CL^pro^) (Gorbalenya et al., 1989). Studies have shown that M^pro^s of difference coronaviruses are highly conserved in terms of both sequences and 3D structures (Xue et al., 2008). These features, together with its functional importance, have made M^pro^ an attractive target for the design of anticoronaviral drugs (Xue et al., 2008; Anand et al., 2003).

To present, these are still no clinically approved antibodies or drugs specific for coronaviruses, which makes it more difficult for curing 2019-nCoV caused diseases and controlling the associated pandemic. With the hope to identify candidate drugs for 2019-nCoV, we adopted a computational approach to screen for available commercial medicines which may function as inhibitors for the M^pro^ of 2019-nCoV.

## Results

A previous attempt to predict drugs for the M^pro^ of SARS-CoV has identified two HIV-1 protease inhibitors, namely lopinavir and ritonavir, as potential candidates, both of which bind to the same target site of M^pro^ (Nukoolkarn et al., 2008). Clinical application of these two drugs on 2019-nCoV patients also appears to be effective, demonstrating the importance of the drug binding site for suppressing 2019-nCoV M^pro^ activity.

To search for other drugs that may inhibit 2019-nCoV M^pro^, we first evaluated the sequence and structural conservation of lopinavir/ritonavir bind site between SARS-CoV and 2019-nCoV. The protein sequences of SARS-CoV M^pro^ and 2019-nCoV M^pro^ are 96% identical (Figure 1a), and the spatial structure of the previously reported lopinavir/ritonavir bind pocket is also conserved between SARS-CoV M^pro^ and 2019-nCoV M^pro^ (Figure 1b). The conserved amino acids Thr24-Asn28 and Asn119 (numbered according to positions in SARS-CoV M^pro^ as the present sequence of 2019-nCoV M^pro^ in GenBank is in polypeptide form) formed the binding pockets for lopinavir/ritonavir in the spatial structure of both SARS-CoV M^pro^ and 2019-nCoV M^pro^, whereas the nearby non-conserved amino acids locate far away from the binding pocket, thus would not affect its structural conservation (Figure 1b). Virtual docking of lopinavir/ritonavir to 2019-nCoV M^pro^ also showed high binding ability to the pocket site (Figure 1c), similar to previous report for SARS-CoV M^pro^ (Nukoolkarn et al., 2008). Amino acids Thr24, Thr26, and Asn119 were predicted to be the key residues for binding the drugs (Figure 1c and Supplementary Figure 1), forming 2 hydrogen bonds with lopinavir and 2 hydrogen bonds with ritonavir, respectively.

**Figure 1.**
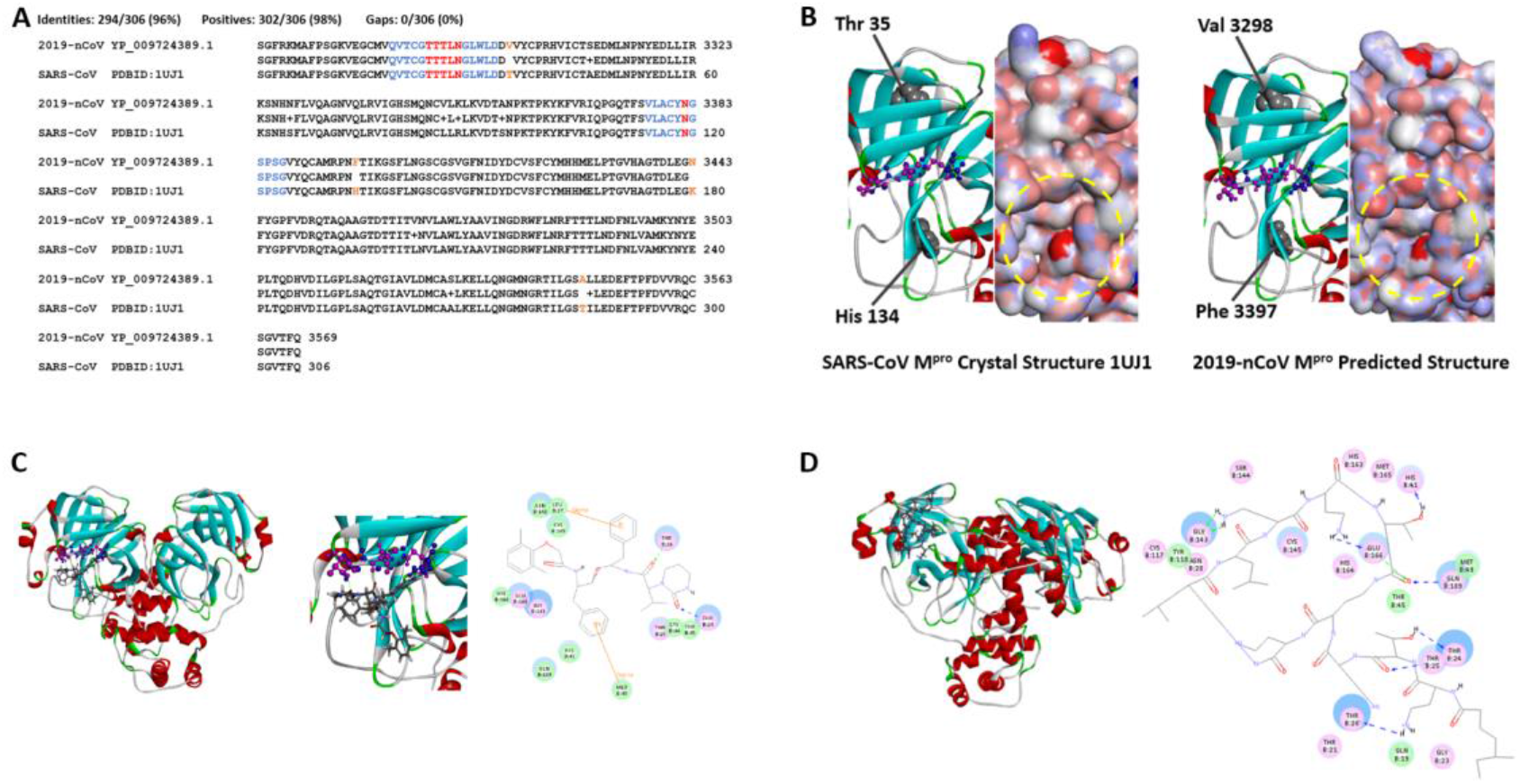
Screen for potential 2019-nCoV M^pro^ inhibitors from commercial medicines. A. Sequence comparison between 2019-nCoV M^pro^ and SARS-Cov M^pro^. Amino acids forming hydrogen bonds with drugs are shown in red, their adjacent 5 amino acids on each side are shown in blue. Mutated amino acids rather than positive substitutions are shown in orange. B. Structural comparison of lopinavir/ritonavir binding pocket in SARS-CoV M^pro^ and 2019-nCoV M^pro^. Ribbon models show the pocket structure of SARS-CoV M^pro^ (left) and 2019-nCoV M^pro^ (right), with α-helixes shown in red and β-sheets in cyan. Residues essential for lopinavir/ritonavir binding are shown in ball-and-stick format, of which Thr24, Thr26, and Asn28 are shown in purple, Thr25, Leu27, and Asn119 are shown in blue. Protein solid surface model is shown to the right of each ribbon model, with the outer rim of the binding pocket marked by dashed yellow cycle. Mutations (marked in orange in panel A) are shown as gray balls, which are apart from the binding pocket. C. Docking model of lopinavir to 2019-nCoV M^pro^. Left, overall docking model of lopinavir to 2019-nCoV M^pro^; middle, enlargement of the lopinavir binding region; right, predicted chemical bonds between lopinavir and key residues of the binding pocket. D. Docking model of colistin to 2019-nCoV M^pro^. Left, overall docking model of colistin to 2019-nCoV M^pro^; right, predicted chemical bonds between colistin and key residues of the binding pocket. In (C) and (D), protein ribbon models are shown with the same diagram as described in (B), drugs are shown as sticks. Hydrogen bonds between drugs and amino acids are shown as dash lines, Pi bonds are shown as orange lines.

Basing on these results, we performed virtual docking to screen for commercial medicines in the DrugBank database that could bind to the above mentioned pocket site of 2019-nCoV M^pro^, and identified 10 candidate clinical medicines (Table 1, Figure 1d and Supplementary Figure 2). These drugs could form hydrogen bonds with one or more residues among Thr24-Asn28 and Asn119, theoretically, are capable to bind to the pocket formed by these amino acids and interfere the function of 2019-nCoV M^pro^.

**Table 1.** Predicted commercial medicines as potential inhibitors for 2019-nCoV M^pro^.

| Name | H-bond counts | Vital residues taking part in H-bond formation | Medical indications according to DrugBank |
| --- | --- | --- | --- |
| Colistin | 9 | THR24, THR25, THR26 | Antibiotic |
| Valrubicin | 7 | THR24, THR25, THR26, ASN28, ASN119 | Anthracycline, antitumor |
| Icatibant | 6 | ASN28, ASN119 | Hereditary angioedema |
| Bepotastine | 5 | THR25, THR26, ASN119 | Rhinitis, urticaria/puritus |
| Epirubicin | 4 | ASN28, ASN119 | Antitumor |
| Epoprostenol | 4 | ASN119 | Vasodilator, platelet aggregation |
| Vapreotide | 3 | THR24, ASN28, ASN119 | Antitumor |
| Aprepitant | 3 | ASN28, ASN119 | Nausea, vomiting, antitumor |
| Caspofungin | 3 | ASN119 | Antifungal |
| Perphenazine | 2 | ASN28, ASN119 | Antipsychotic |

In summary, basing on the structural information of clinical effective medicines for 2019-nCoV, we have predicted a list of commercial medicines which may function as inhibitors for 2019-nCoV by targeting its main protease M^pro^. Compared to lopinavir/ritonavir, most of these predicted drugs could form more hydrogen bounds with 2019-nCoV M^pro^, thus may have higher mutation tolerance than lopinavir/ritonavir. The binding pockets of these drugs on M^pro^ are conserved between SARS-CoV M^pro^ and 2019-nCoV M^pro^, indicating the potential of these drugs to function as inhibitors for other coronaviruses with similar M^pro^ binding sites and pocket structures.

## Acknowledgements

This work is supported by grants from CAS Advance Research Programs (QYZDJ-SSW-SMC015) and National Natural Science Foundation of China (91940304 and 81790622) to X.-J. W.

## Methods

### Sequence Resource

The protein sequences of SARS-CoV M^pro^ (Accession: 1UK3_A) and 2019-nCoV polyprotein orf1ab (Accession: YP_009724389.1) were downloaded from GenBank (http://www.ncbi.nlm.nih.gov). The protein sequence of 2019-nCoV M^pro^ was determined by aligning the SARS-CoV M^pro^ sequence to 2019-nCoV polyprotein orf1ab using BLAST (Altschul et al., 1990), the best aligned region in 2019-nCoV orf1ab to SARS-CoV M^pro^ was selected as 2019-nCoV M^pro^.

### Structure Modeling

Crystal structure of SARS-CoV M^pro^ (PDB ID: 1UJ1) was downloaded from Protein Data Bank (PDB, http://www.rcsb.org) (Burley et al., 2019). Structure of 2019-nCoV M^pro^ was predicted by Modeller algorithm (Webb and Sali, 2016) using the structure of SARS-CoV M^pro^ as template. Structural details were visualized with the Visualizer function of Discovery Studio 3.5 (Accelrys Software Inc).

### Candidate Drug Screen

Molecular structures of commercial available medicines were downloaded from the DrugBank database (http://www.drugbank.ca) (Wishart et al., 2018). The original indications of medicines were collected according to DrugBank descriptions. Virtual screening of medicines with binding potential to the pocket site of 2019-nCoV M^pro^ was performed using the Libdock algorithm of Discovery Studio 3.5 (Accelrys Software Inc). The pocket site of 2019-nCoV M^pro^ was identified by homology comparison to a previous published work (Nukoolkarn et al., 2008). Spatial conformations of medicines were generated using CAESAR algorithm of Discovery Studio 3.5 using default parameters. Maximal hits of the docking process was set to 50.

---

**Supplementary Figure 1.**
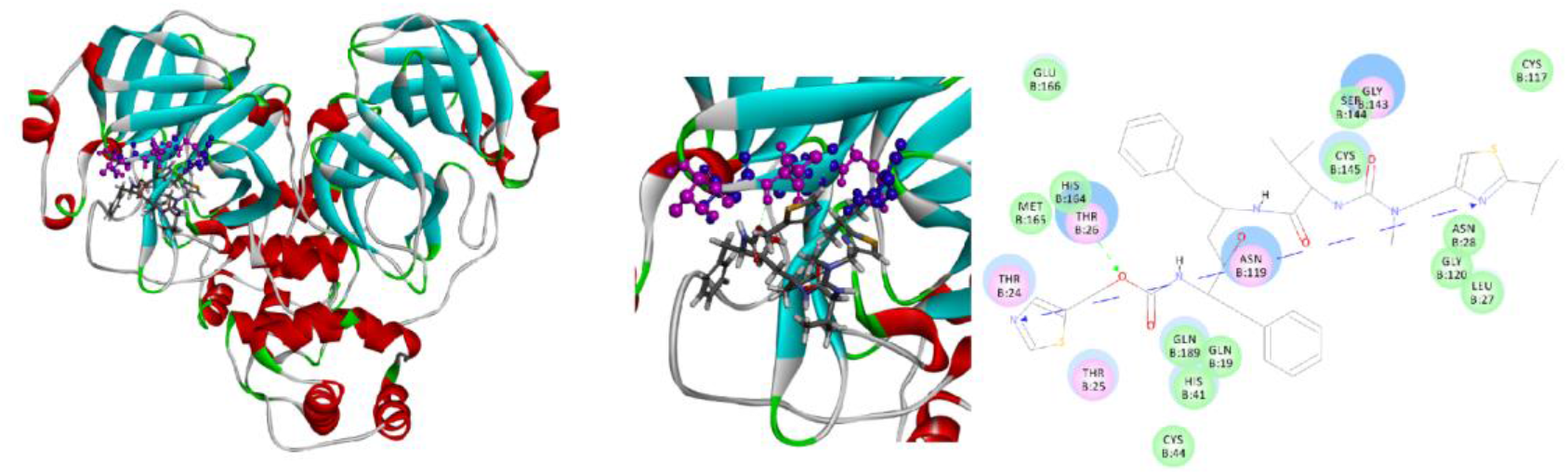
Docking model of ritonavir to 2019-nCoV M^pro^. Left, overall docking model of ritonavir to 2019-nCoV M^pro^; middle, enlargement of the ritonavir binding region; right, predicted chemical bonds between ritonavir and key residues of the binding pocket. Ribbon model shows the pocket structure of 2019-nCoV M^pro^ (right), with α-helixes shown in red and β-sheets in cyan. Residues essential for ritonavir binding are shown in ball-and-stick format, of which Thr24, Thr26, and Asn28 are shown in purple, Thr25, Leu27, and Asn119 are shown in blue. Ritonavir is shown as sticks. Hydrogen bonds between drugs and amino acids are shown as dash lines, Pi bonds are shown as orange lines.

**Supplementary Figure 2.**
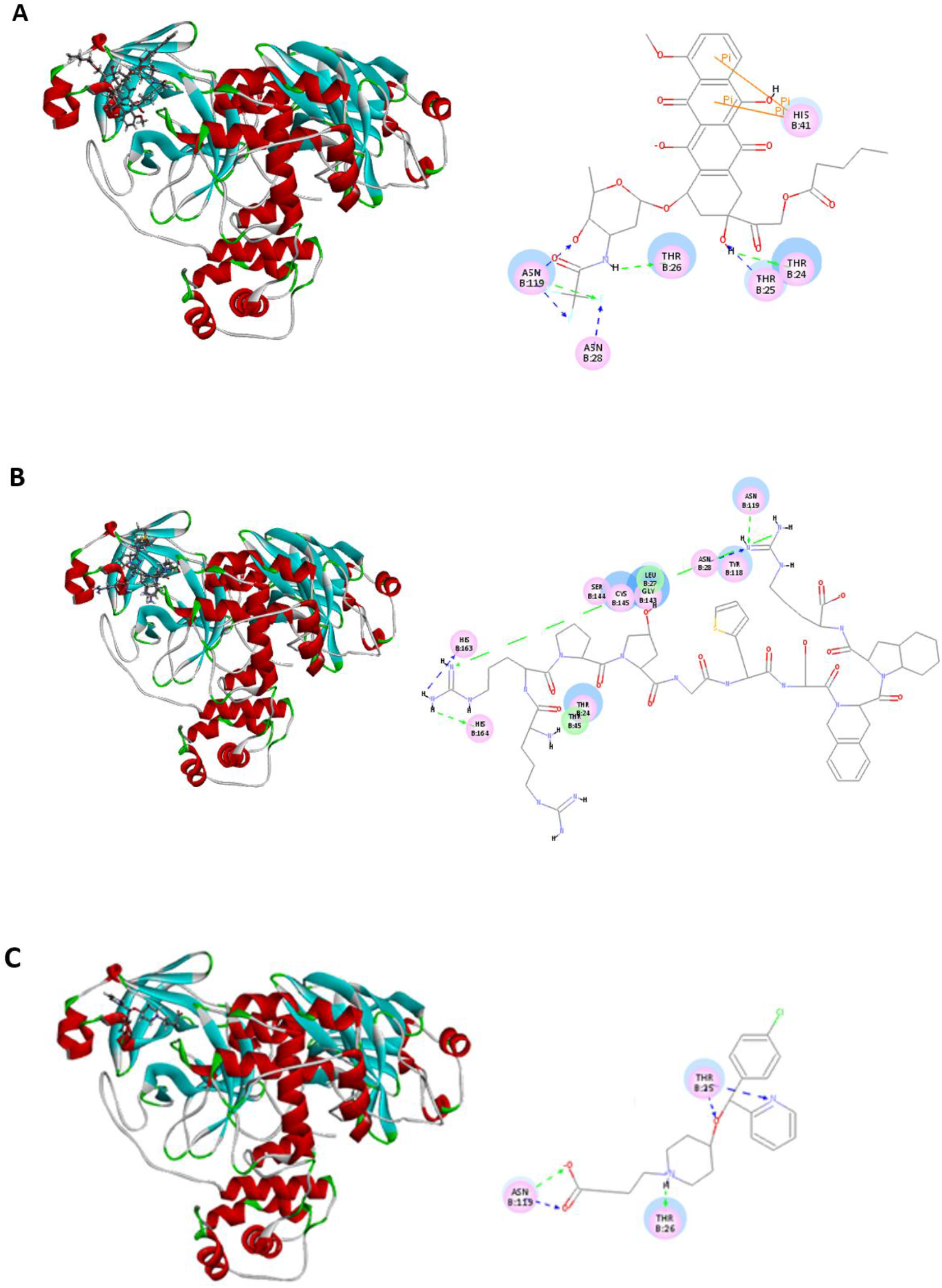
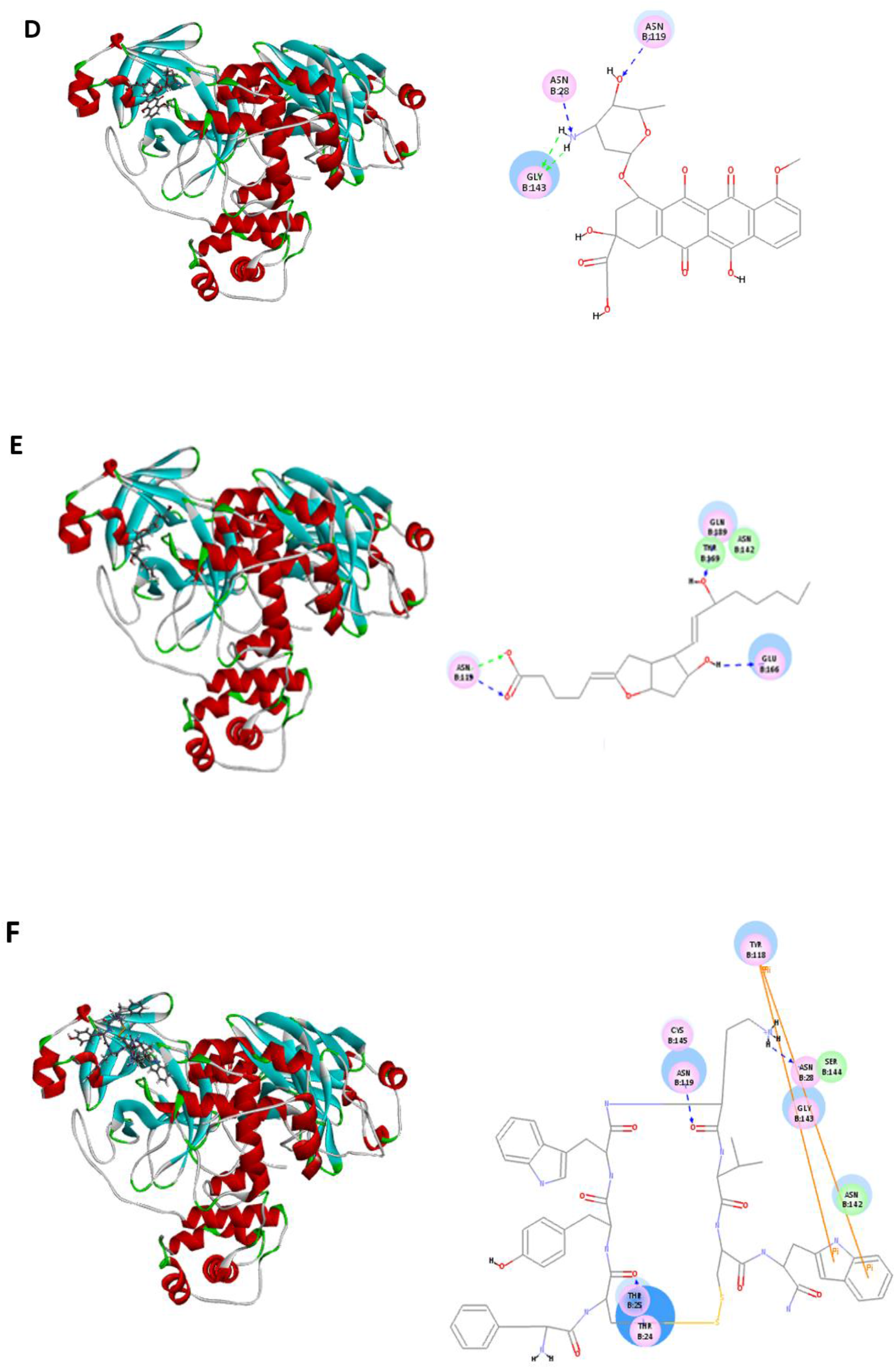
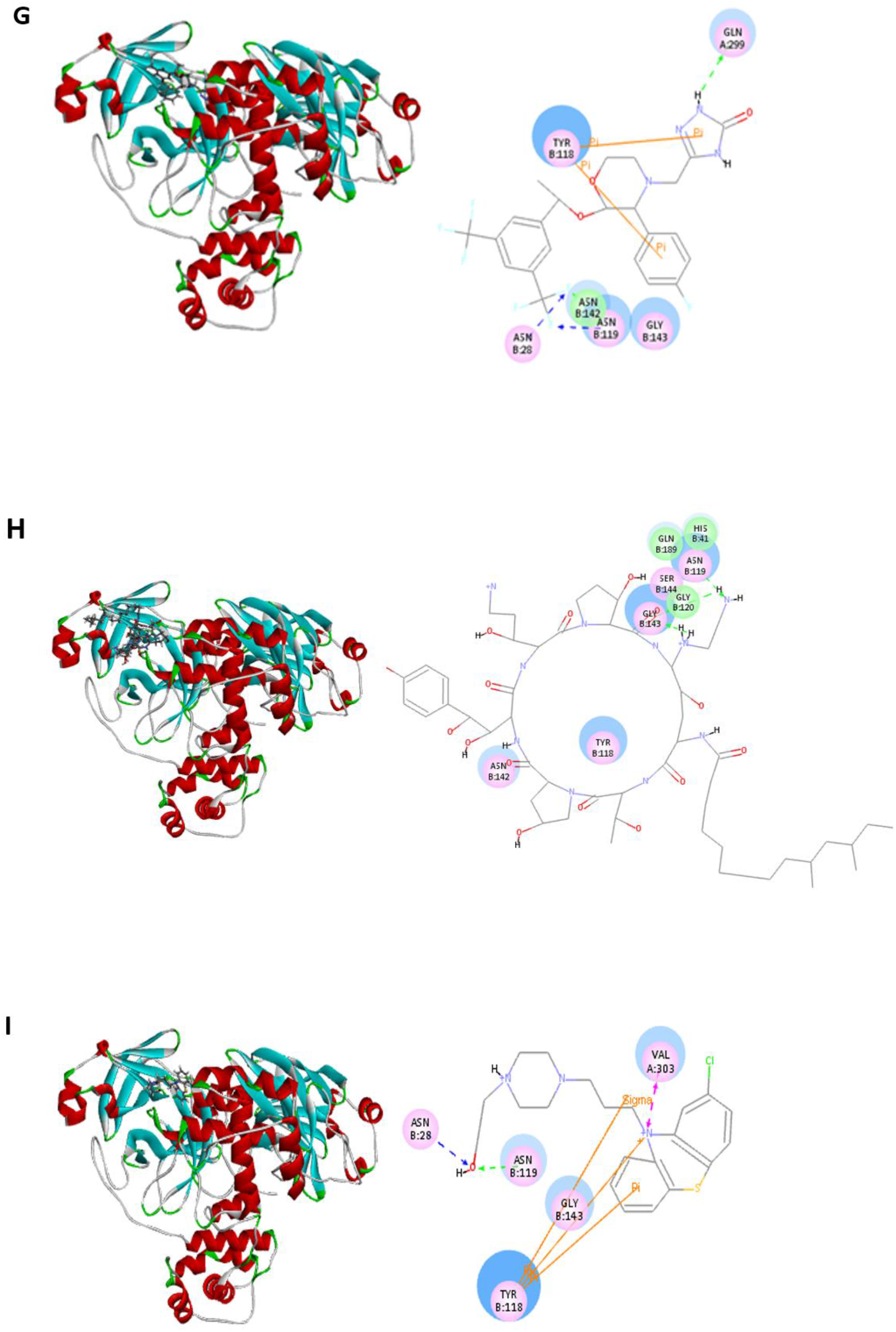
Docking model of candidate inhibitors drugs to 2019-nCoV Mpro. A. Valrubicin; B. Icatibant; C. Bepotastine; D. Epirubicin; E. Epoprostenol; F. Vapreotide; G. Aprepitant; H. Caspofungin; I. Perphenazine. Legends are the same as described in Figure 1d.

## References

Anand, K., Ziebuhr, J., Wadhwani, P., Mesters, J.R., and Hilgenfeld, R., 2003. Coronavirus main proteinase (3CLpro) structure: basis for design of anti-SARS drugs. Science. 300, 1763–1767.

Cui, J., Li, F., and Shi, Z.L., 2019. Origin and evolution of pathogenic coronaviruses. Nat. Rev. Microbiol. 17, 181–192.

Dhama, K., Pawaiya, R., Chakraborty, S., Tiwari, R., M, S., and Verma, A., 2014. Coronavirus Infection in Equines: A Review. Asian J. Anim. Vet. Adv. 9, 164–176.

Gorbalenya, A.E., Donchenko, A.P., Blinov, V.M., and Koonin, E.V., 1989. Cysteine proteases of positive strand RNA viruses and chymotrypsin-like serine proteases. A distinct protein superfamily with a common structural fold. FEBS Lett. 243, 103–114.

Hui, D.S., E, I.A., Madani, T.A., Ntoumi, F., Kock, R., Dar, O., Ippolito, G., McHugh, T.D., Memish, Z.A., Drosten, C., Zumla, A., and Petersen, E., 2020. The continuing 2019-nCoV epidemic threat of novel coronaviruses to global health - The latest 2019 novel coronavirus outbreak in Wuhan, China. Int. J. Infect Dis. 91, 264–266.

Lai, M.M.C., Holmes, K.V., 2001. Coronaviridae: the viruses and their replication, in: Knipe, D.M. and Howley, P.M. (Eds.), Fields virology. Lippincott Williams & Wilkins, Philadelphia, pp. 1163–1179.

Lu, H., Stratton, C.W., and Tang, Y.W., 2020. Outbreak of Pneumonia of Unknown Etiology in Wuhan China: the Mystery and the Miracle. J. Med. Virol. DOI: 10.1002/jmv.25678.

Masters, P.S., Perlman, S., 2013. Coronaviridae, in: Knipe, D.M. and Howley, P.M. (Eds.), Fields virology. Lippincott Williams & Wilkins, Philadelphia, pp. 825–858.

Nukoolkarn, V., Lee, V.S., Malaisree, M., Aruksakulwong, O., and Hannongbua, S., 2008. Molecular dynamic simulations analysis of ritonavir and lopinavir as SARS-CoV 3CL(pro) inhibitors. J. Theor. Biol. 254, 861–867.

Regenmortel, M.H.V. van, Fauquet, C.M., Bishop, D.H.L., Carstens, E.B., Estes, M. K., Lemon, S.M., Maniloff, J., Mayo, M.A., McGeoch, D.J., Pringle, C.R., Wickner, R.B., 2000. Coronaviridae, in: Regenmortel, M.H.V. van, Fauquet, C.M., Bishop, D.H.L., et al. (Eds.), Virus taxonomy: Classification and nomenclature of viruses. Seventh report of the International Committee on Taxonomy of Viruses. Academic Press, San Diego, pp. 835–849.

Schoeman, D., and Fielding, B.C., 2019. Coronavirus envelope protein: current knowledge. Virol. J. 16, 69.

Xue, X., Yu, H., Yang, H., Xue, F., Wu, Z., Shen, W., Li, J., Zhou, Z., Ding, Y., Zhao, Q., Zhang, X.C., Liao, M., Bartlam, M., and Rao, Z., 2008. Structures of two coronavirus main proteases: implications for substrate binding and antiviral drug design. J. Virol. 82, 2515–2527.

Zhu, N., Zhang, D., Wang, W., Li, X., Yang, B., Song, J., Zhao, X., Huang, B., Shi, W., Lu, R., Niu, P., Zhan, F., Ma, X., Wang, D., Xu, W., Wu, G., Gao, G.F., and Tan, W., 2020. A Novel Coronavirus from Patients with Pneumonia in China, 2019. N. Engl. J. Med. DOI: 10.1056/NEJMoa2001017.

## References

Altschul, S.F., Gish, W., Miller, W., Myers, E.W., and Lipman, D.J., 1990. Basic local alignment search tool. J. Mol. Biol. 215, 403–410.

Burley, S.K., Berman, H.M., Bhikadiya, C., Bi, C., Chen, L., Di Costanzo, L., Christie, C., Dalenberg, K., Duarte, J.M., Dutta, S., Feng, Z., Ghosh, S., Goodsell, D.S., Green, R.K., Guranovic, V., Guzenko, D., Hudson, B.P., Kalro, T., Liang, Y., Lowe, R., Namkoong, H., Peisach, E., Periskova, I., Prlic, A., Randle, C., Rose, A., Rose, P., Sala, R., Sekharan, M., Shao, C., Tan, L., Tao, Y.P., Valasatava, Y., Voigt, M., Westbrook, J., Woo, J., Yang, H., Young, J., Zhuravleva, M., and Zardecki, C., 2019. RCSB Protein Data Bank: biological macromolecular structures enabling research and education in fundamental biology, biomedicine, biotechnology and energy. Nucleic Acids Res. 47, D464–D474.

Wishart, D.S., Feunang, Y.D., Guo, A.C., Lo, E.J., Marcu, A., Grant, J.R., Sajed, T., Johnson, D., Li, C., Sayeeda, Z., Assempour, N., Iynkkaran, I., Liu, Y., Maciejewski, A., Gale, N., Wilson, A., Chin, L., Cummings, R., Le, D., Pon, A., Knox, C., and Wilson, M., 2018. DrugBank 5.0: a major update to the DrugBank database for 2018. Nucleic Acids Res. 46, D1074–D1082.

